# Facing lethal temperatures: heat shock response in desert and temperate ants

**DOI:** 10.1101/2022.11.12.516273

**Authors:** Natalia de Souza Araujo, Rémy Perez, Quentin Willot, Matthieu Defrance, Serge Aron

**Affiliations:** Department of Evolutionary Biology & Ecology, Université Libre de Bruxelles. Avenue F.D. Roosevelt, 50. B-1050 Brussels, Belgium; Interuniversity Institute of Bioinformatics in Brussels, Université Libre de Bruxelles. B-1050 Brussels, Belgium; Zoophysiology, Department of Biology, Aarhus University, 8000, Aarhus-C, Denmark

## Abstract

Several genera of desert ants have adapted to endure prolonged exposure to high temperatures. The study of these ants is essential to unravel how species respond and adapt to thermal stress. We investigated the thermal tolerance and the transcriptomic heat stress response of three desert ant genera (*Cataglyphis, Melophorus* and *Ocymyrmex*) and two temperate genera (*Formica* and *Myrmica*) to explore convergent and specific adaptations. We found a variable transcriptomic response among desert species exposed to similar levels of physiological heat-stress: *Cataglyphis holgerseni* and *Melophorus bagoti* differentially regulated very few transcripts, 0.12% (54/44,525) and 0.14% (53/38,726) respectively, while *Cataglyphis bombycina* and *Ocymyrmex robustior* showed greater expression alterations affecting 0.6% (253/41,912) and 1.53% (698/45,701) of their transcriptomes, respectively. These two responsive mechanisms – reactive and constitutive – were related to desert species thermal tolerance survival pattern and convergently evolved in distinct desert ant genera. By comparison, the two temperate species differentially expressed thousands of transcripts more than desert ants in response to heat stress (affecting 8% and 12,71% of *F. fusca* and *Myr. sabuleti* transcriptomes), suggesting that keeping restrained gene expression is an important adaptation in heat adapted species. Finally, we found a significant overlap of the molecular pathways activated in response to heat-stress in temperate and desert species, and our data revealed that larger gene expression responses also affected a greater number of taxonomically restricted genes. These results suggest that the molecular processes involved in heat-stress response are mostly evolutionary conserved in ants, but new genes may also play a role.

## Introduction

Global climate changes affect species distribution, adaptation, and survival [1]. Ectotherm species are particularly highly sensitivity to temperature increase, and even moderate warmer temperatures may lead to a disproportionate increased heat failure [2]. To establish and improve predictive models and management plans to control biodiversity loss, it is therefore primordial to understand the effects of thermal stress in different organisms [2,3]. A major effect of severe heat stress is the denaturation of the three-dimensional structure of macromolecules such as proteins and cell membranes, leading to function loss, metabolic failure and, ultimately, cellular and organism death [4,5]. To face heat-stress, the cellular machinery may activate multiple pathways increasing the production of proteins to limit or repair cell damage [6]. The best-known examples of proteins involved in this response are the heat-shock proteins (HSP). These proteins perform several functions essential to survival; they ensure correct refolding of denaturated proteins, prevent harmful protein aggregation (due to the association of misfolded proteins), and participate in the elimination of protein aggregates [7]. Other pathways triggered by high temperature include that of unsaturated longer fatty acids and sterols accumulation, whose increased production helps to keep the fluidity of the cell matrix and of the membrane [8]. Likewise, the production of antioxidants and detoxification enzymes, like super oxide dismutase (SOD) or glutathione peroxidase (GPx), helps reducing the accumulation of reactive oxygen species (ROS) [9]. Under heat stress, these molecular pathways interact composing an intricate multi-level response that, despite its complexity, must be rapidly activated to optimize survival [10]. This response is expected to be particularly important in small ectotherms – like insects – whose body temperature is highly correlated to that of their local environment [11].

Hot deserts are amongst the most stressful habitats on Earth. Organisms inhabiting these regions must cope with extremely high temperatures and low humidity [12]. These conditions have led most desert animals to adopt nocturnal or bimodal activity patterns, exploiting the cooler dawn and twilight hours to avoid the scorching heat [13,14]. Amazingly, several ant species display the exact opposite behaviour; they only leave the nest during the warmest hours of the day when air and ground temperature are close to their own lethal thermal limits [15–17]. Such unusual activity pattern allows these insects to forage for heat-struck arthropods while reducing competition and predation pressures from other desert organisms. This so called “thermal scavenging” behaviour has appeared at least three distinct times in phylogeographically distant heat adapted ant genera: (A) within the *Cataglyphis* in the deserts of the northern hemisphere, (B) in some *Melophorus* species from the Australian outback, and (C) among the *Ocymyrmex* who inhabit deserts of sub-tropical Africa (Fig 1; [18–20]). Workers of these genera exhibit similar behavioural and morphological adaptations likely resulting from a convergent evolutionary process in response to their peculiar thermal niche [21]. They can withstand body temperature exceeding 45°C during a long period of time [22–24]; their legs are considerably longer relatively to their body size, which maximizes the distance between their body and the burning ground and allows higher running speed reducing foraging time and enhancing forced convective cooling [25,26]; and they periodically exploit thermal refuges in shaded areas or ground elevated places to benefit from cooler spots [20]. Recently, our group has characterized the transcriptomic heat shock response of six *Cataglyphis* species from different habitats and thermal regimes [10,27]. We found that the *Cataglyphis* heat stress response builds upon the co-expression of gene clusters mainly involved in proteome stability, elimination of toxic residues, DNA/RNA metabolism and the maintenance of cell integrity through membrane modification and cytoskeletal rearrangements. Whether such molecular adaptations/pathways to cope with heat stress are genera-specific or convergently evolved in different desert ants remains however unknown.

**Fig 1.**
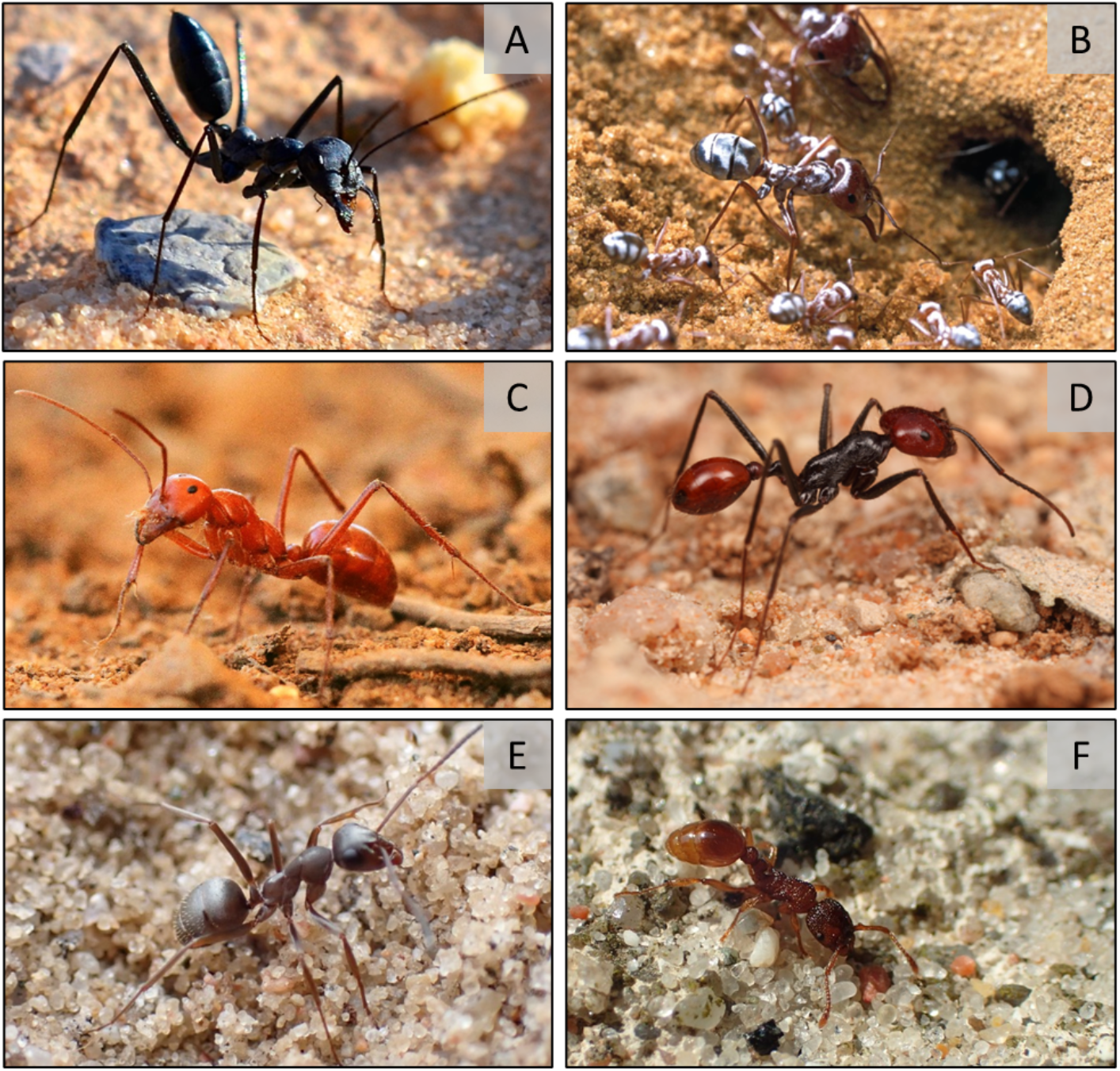
Workers of the ant species:(A) *Cataglyphis holgerseni*, (B) *C. bombycina*, (C) *Melophorus bagoti*, (D) *Ocymyrmex sp*., (E) *Formica fusca*, and (F) *Myrmica sabuleti*. With permission: (A) A. Kuhn, (B, E, F) Q. Willot, (C) A. Wystrach, (D) R. Duncan.

Herein, we investigated the thermal tolerance and the transcriptomic response to heat stress of four species of desert ants belonging to the genera *Cataglyphis, Melophorus* and *Ocymyrmex* (Fig 1). Additionally, to explore convergent and specific adaptations to heat-stress, we compared the heat-shock response found in these desert ants with that of related outgroup temperate species from the Formicinae (same subfamily as *Cataglyphis* and *Melophorus*) and Myrmicinae (same subfamily as *Ocymyrmex*). We first measured their heat tolerance by assessing the survival rate of workers at increasing temperatures and established the species thermal limit. Then, we identified genes involved in the heat-shock response by quantifying the differentially regulated transcripts between heat stressed and control workers. Finally, we performed cross-species comparisons using differential gene expression and survival results to identify whether any convergence could be observed in the heat shock response of desert species. We show that desert ants benefit from two main molecular strategies to deal with high temperatures. The first is a constitutive high expression of heat stress genes; this response is associated with individuals’ survival decreasing rapidly after reaching their thermal limit. The second, a reactive gene expression regulation in response to increasing temperatures that was associated with a wider variation in individual survival capacity above species thermal limit. Additionally, unlike temperate species, desert ants maintain strong gene expression control even at considerably higher temperatures, indicating this is a key adaptation to withstand heat stress.

## Results

### Desert ants show two survival patterns aligned to two distinct gene expression regulation responses under heat stress

We estimated ant heat tolerance through heat-stress assays in which workers were exposed to constant temperatures (39°C, 41°C, 43°C, 45°C, 47°C, 49°C or 51°C). In total six groups of 10 randomly selected workers (totalizing 60 individuals) were tested per species, and after 3 hours of exposure to the tested temperature the per cent survival of each group was recorded. The median lethal temperature after 3h exposure (LT50_3h) differed significantly among species (Fig S1). It was 39.91°C (CI 95%: ±0.16) for *Myr. sabuleti;* 42°C (±0.17) for *F. fusca;* 44.83°C (±0.18) for *C. bombycina;* 46°C (±0.16) for *C. holgerseni;* 46.8°C (±0.16) for *O. robustior;* and 49.83°C (±0.15) for *Mel. bagoti*. Consistently, the temperature below which the survival rate was significantly different from 100% after 3h exposure (upper thermal limit, UTL_3h) was also the lowest for the two temperate species, *Myr. sabuleti* and *F. fusca*, at 39°C and 41°C respectively. *Mel. bagoti* was the most heat tolerant ant species with 95% of workers surviving at 49°C for 3 hours. The other ants showed intermediated UTL_3h values; *C. holgerseni* had a UTL_3h of 45°C, and both *C. bombycina* and *O. robustior* an UTL_3h of 43°C. For all species, the survival ratio markedly dropped from about 100% to 0% within a two-degree temperature increase interval (Fig 2), except for the Sahara ant *C. bombycina* and the Namib ant *O. robustior*, whose survival gradually decreased within a wider temperature range of six degrees.

**Fig 2.**
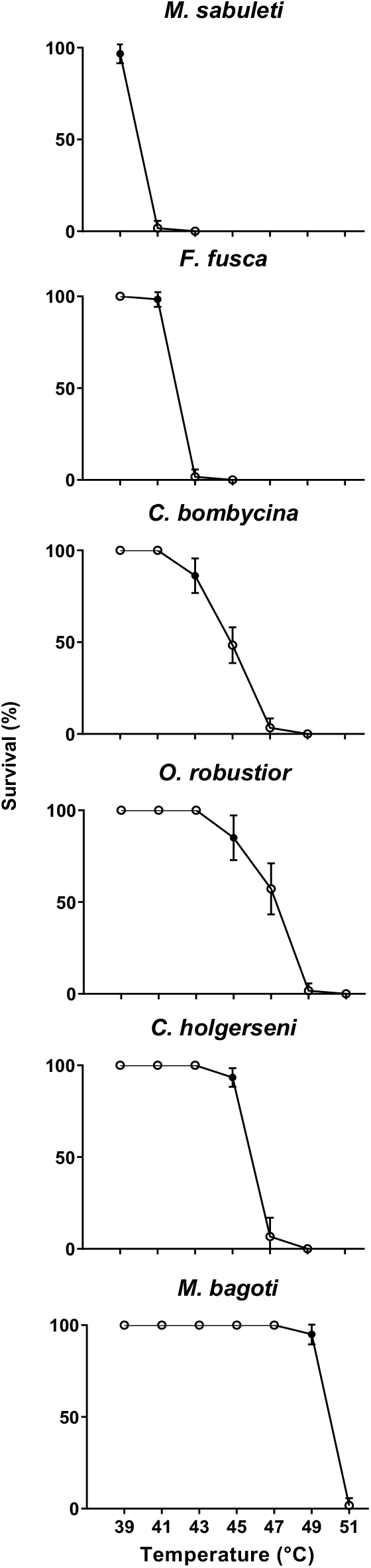
Percent survival of ant workers during heat-stress assays. The ants (60 workers from 4-6 colonies per species) were exposed to increasing temperatures (39°C, 41°C, 43°C, 45°C, 47°C, 49°C or 51°C) for 3 hours; their percent survival was then measured. Workers were categorized as moribund when reaching muscular paralysis. For each species, UTL is indicated by a filled circle (•).

Reference transcriptomes were assembled using RNASeq data from a pool of workers (Heat Stressed – HS – and Non Heat Stressed – NHS), eggs, and pupae. For species with a reference genome available (*i.e*., the *Cataglyphis* species and *F. exsecta*) we generated a reference guided and a *de novo* assembly; for *O. robustior, Mel. bagoti* and *Myr. Sabuleti*, only a *de novo* assembly was performed. In all cases, we clustered similar transcripts using CD-Hit [28], Corset [29] and Lace [30], and removed non protein coding transcripts. Reference transcriptome sizes ranged from 38,726 in *Mel. bagoti* to 84,784 in *Myr. sabuleti* (Fig 3B, major quality parameters are given in Table S2). The cause for such variation in transcriptome size among species is unknown; it might stem from the presence of multiple isoforms, the nature of the transcripts (CG or AT rich) or the quality of the reference genome used. Assembly strategy (with or without the reference genome) did not relate directly with the total number of transcripts, as species with a reference genome did not necessarily have less transcripts. Instead, transcripts were likely more complete – with higher N50 values and more complete BUSCOs (Table S2) – when using the reference genome.

**Fig 3.**
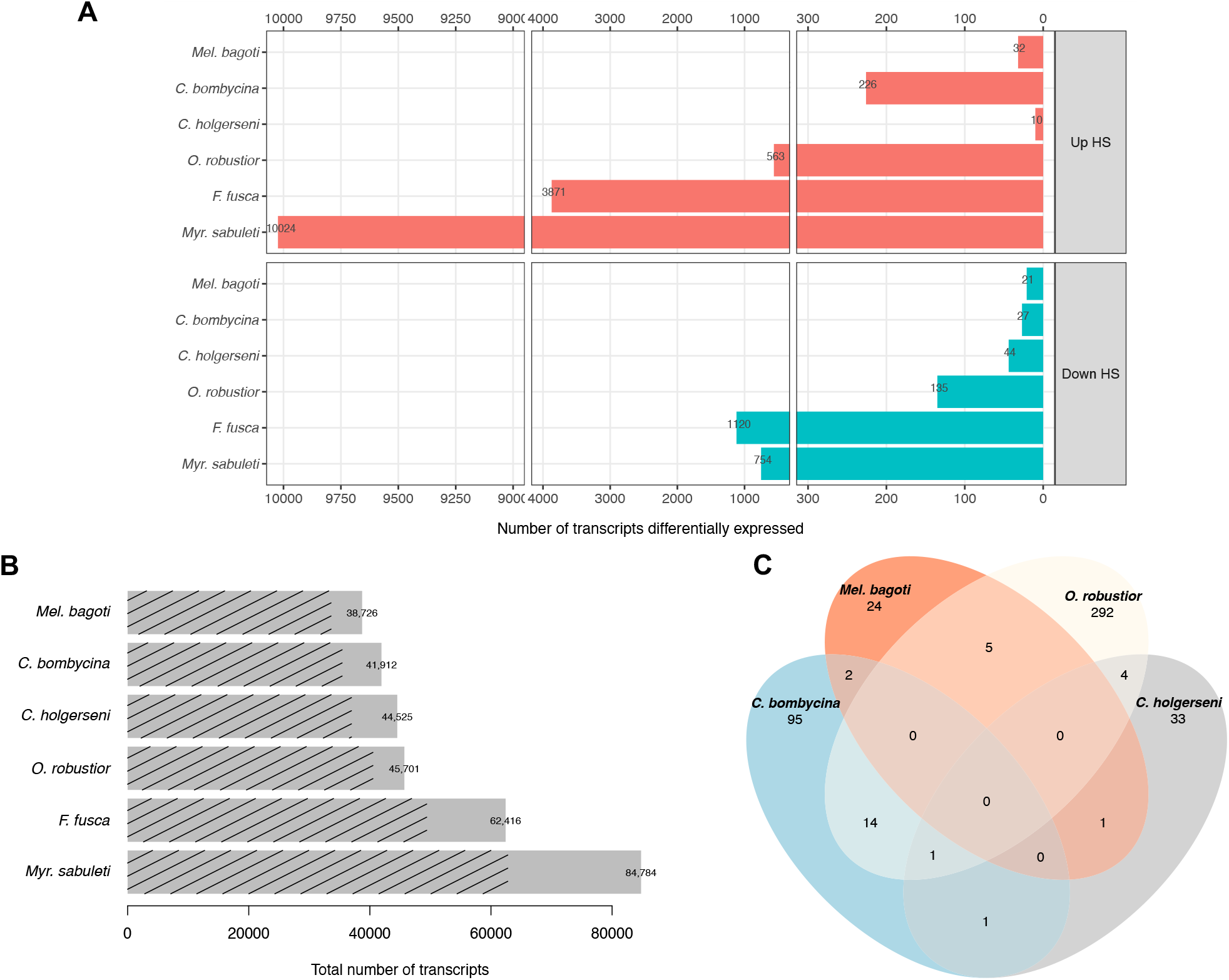
Transcriptomic analyses summary. **A** – Number of differentially expressed transcripts between heat stressed and non-heat stressed conditions for each species tested. Up HS transcripts refers to transcripts up regulated in the heat stressed samples when compared to the non-heat stressed samples, while Down HS refers to transcripts down regulated in heat stressed samples (i.e., up regulated in non-heat stressed samples in comparison to heat-stressed ones). **B** – Reference transcriptome sizes. Dashed lines show the proportion of transcripts with at least one significant blast hit. **C** – Number of transcripts differentially expressed and annotated in common among desert species. Only the overlap between *C. bombycina* and *O. robustior* was significant at p < 0.01 random sampling test. For the sake of clarity, overlapping results are shown only for desert species. All species comparisons are given in Table S3.

Differentially expressed transcripts (DET) were identified using four replicates per treatment; in the HS treatment, workers were exposed to the species upper thermal limit temperature for 3h; in the NHS treatment, workers were maintained for 3h hours at 25°C (desert species) or 18°C (temperate species). Ten worker whole bodies were pooled for RNA sequencing of each sample replicate. The number of transcripts differentially expressed between the HS and NHS conditions ranged from 53 in *Mel. bagoti* to 10,778 in *Myr. sabuleti* (Fig 3A; complete lists of DET in each species and their annotations are provided in Supplementary Files 2–13). Based on the range of differential gene expression observed across the species, we grouped their responses into three responsive profiles that were not phylogenetically conserved (Fig 4): the first comprised *Mel. bagoti* and *C. holgerseni*, both species showing weak heat stress expression responses with only 53 DET in 38,726 transcripts (0.14%) and 54 DET in 44,525 transcripts (0.12%), respectively; the second included *C. bombycina* and *O. robustior*, which had a medium heat stress expression response of 253 DET in 41,912 transcripts (0.60%) and 698 DET in 45,701 transcripts (1.53%), respectively; the third group contained *F. fusca* and *Myr. sabuleti* which had strong heat stress responses and differentially expressed 4,991 in 62,416 transcripts (8%) and 10,778 in 84,784 transcripts (12,71%), respectively. We found a highly significant correlation between transcriptome size and the number of transcripts differentially expressed in response to heat stress across all species (*Pearson R* = 0.99, *p* = 6e-05, Fig S2). However, when only desert species were considered, this correlation was lost (*Pearson R* = 0.63, *p* = 0.37, Fig S2). This suggests that the difference in the number of DET between desert and temperate species is at least partly due to differences in their transcriptome sizes. Nevertheless, the difference between transcriptome sizes and the number of DET were of distinct magnitudes; temperate species showed a minimum seven-fold increase in gene expression activity during assays in comparison to desert ants (Fig 3A), while the maximum transcriptome size difference was only one-fold greater (Fig 3B).

**Fig 4.**
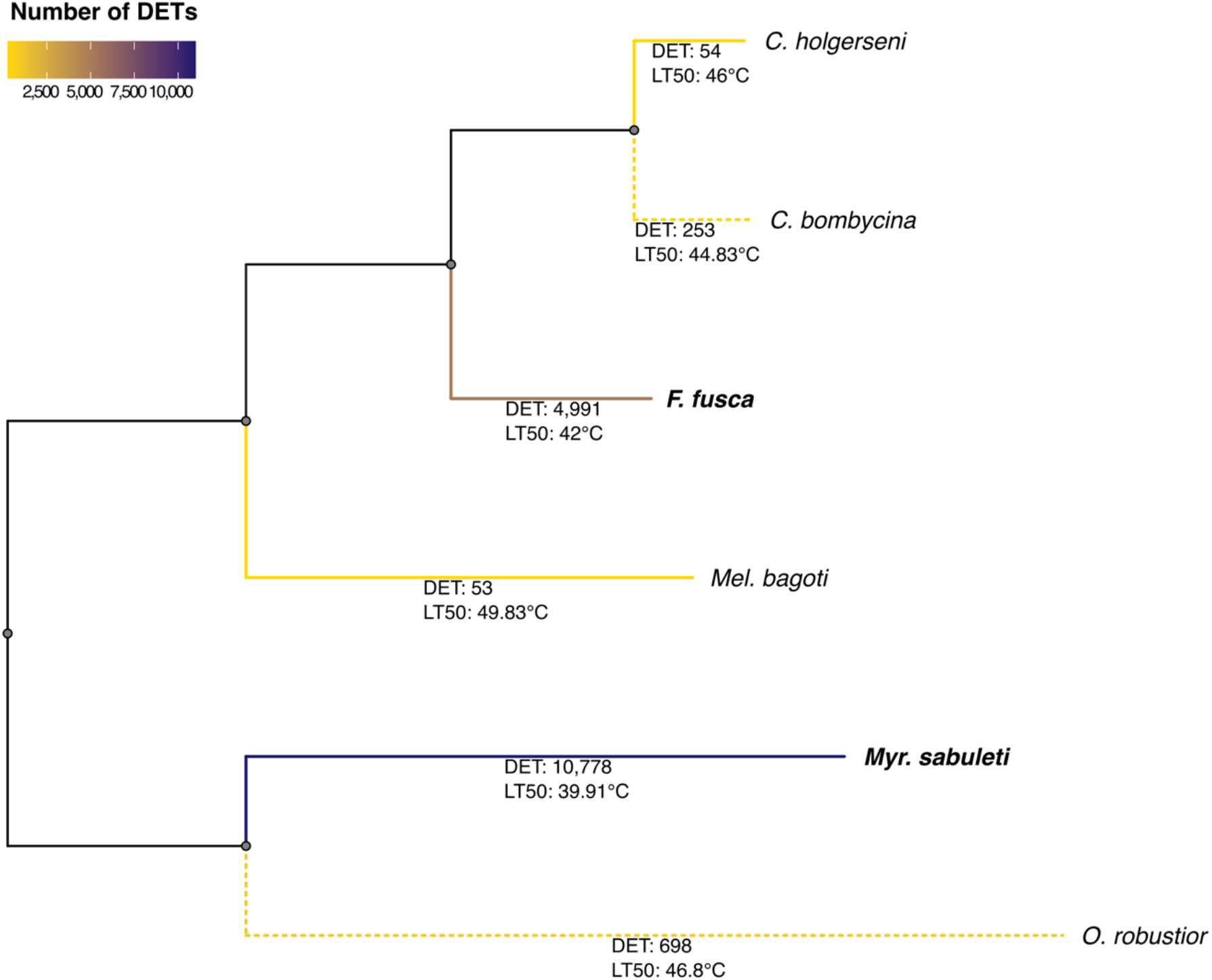
Phylogenetic tree of the studied ant species estimated using orthologous proteins. Branch colours corresponds to the number of differentially expressed transcripts (DET) identified for that species between heat stressed and non-heat stressed samples. The exact number of DET is reported bellow each species branch along with the LT50. Grey lines represent internal nodes with no values. Dotted lines indicate species showing gradual decrease in survival ratio within a six-degree temperature interval while straight lines refer to species whose survival decline from 100 to 0% within a smaller temperature range of two degrees. Ant species from temperate areas are shown in bold.

In conclusion, desert ants displayed a more restrained gene expression regulation in response to thermal stress than temperate species, and transcriptome size variation alone cannot account for all the observed difference. Furthermore, amongst desert ants, wider gene expression alterations in response to heat stress were linked to larger variation in workers’ survival: species with a more intense expression response (*i.e., C. bombycina* and *O. robustior*; Fig 4) displayed a more gradual decrease in survival ratio (Fig 2) – dropping from 100% to 0% survival within a six-degree temperature interval – while species with weak heat stress expression response (*i.e*., *C. holgerseni* and *Mel. bagoti*) showed a marked reduction in survival within a smaller temperature range of two degrees. In temperate species, survival also decreased rapidly within a two-degree temperature increase; however, in contrast to desert ants, many genes where differentially expressed.

### Fewer gene expression alterations in response to heat stems from high constitutive levels of heat stress related genes in desert ants

To further investigate whether the different expression patterns found among desert ants were due to a lack of expression of heat stress related genes, or instead, to a distinct timing regulation of these genes, we focussed on the expression pattern of admittedly known heat stress related genes – the heat shock proteins 70 and 90 (HSP70 and HSP90) – and inspected the expression of their related transcripts (in TMM) in HS and NHS replicates. The expression plots of these transcripts in all species are reported in supplementary files S14 to S19. Although expression counts of desert ants HSP70 and HSP90 transcripts under heat stress and non-heat stress conditions were only significantly differentially regulated in *C. bombycina* and *O. robustior*, these genes were still expressed at a high level in *C. holgerseni* and *Mel. bagoti* (Fig 5). In the two latter species, however, expression of HSP70 and HSP90 in the control group (NHS ants) were already elevated, which explains the lack of significant differences between the two groups. This indicates that desert species with fewer gene expression alterations (*i.e*., *C. holgerseni* and *Mel. bagoti*) have a constitutive up regulation of HSP70 and HSP90 and do not significantly change their expression levels when exposed to increasing temperatures, unlike *C. bombycina* and *O. robustior*.

**Fig 5.**
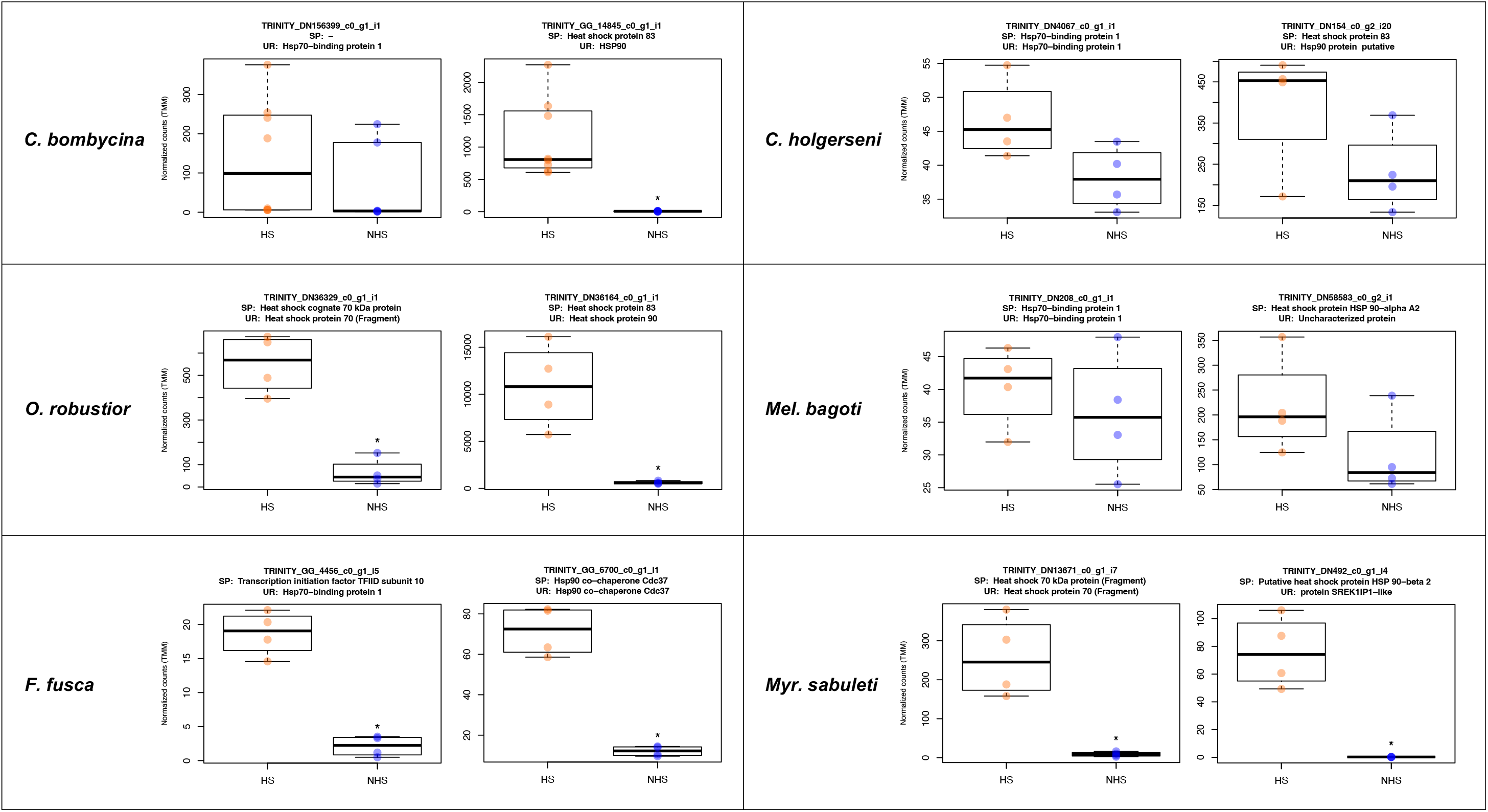
Expression of heat stress proteins 70 and 90 related transcripts in heat-stressed and non-heat stressed treatments. Expression levels shown as TMM normalized read counts. Transcripts shown represent one related transcript from each HSP that was either differentially expressed or had highest expression in the referred species. HS – samples heat stressed; NHS – non-heat stressed control samples; SP – UniProtKB/Swiss-Prot annotation; UR – UniRef90 database annotation. * Indicates the transcript is significantly differentially regulated between treatment conditions (log_2_-fold change ≥ 2 between treatments and FDR ≤1e-3).

These results support the hypothesis that the reduced number of differentially expressed genes under heat stress in *C. holgerseni* and *Mel. bagoti* stems from a constitutive upregulation of heat stress related genes. Therefore, the distinct expression patterns observed in desert ants would reflect a difference in timing regulation of heat stress related genes with *C. holgerseni* and *Mel. bagoti* constantly keeping a high expression of these genes, while *C. bombycina* and *O. robustior* alter their expression in response to heat stress.

### Molecular pathways responsive to heat stress in desert ants with greater gene expression changes are also activated in temperate species

Among desert ants, only *C. bombycina* and *O. robustior* had a significant number of DET in common (Fig 3C). When including the ant species from temperate regions in the comparisons, however, there was a significant number of DET in common among all species except *C. holgerseni* and *Mel. bagoti* (Table S3). Species pair comparisons containing the list of genes found to be differentially expressed in common across species follows in supplementary file S20. Similarly, from the GO analyses (Table S4, Fig S3) we found that terms associated with protein folding (“Protein folding”, “Protein refolding” and *“de novo* protein folding”, respectively GO:0006457, GO:0042026 and GO:0006458) and cellular response to heat (GO:0034605) were commonly enriched in *O. robustior, Myr. sabuleti* and *F. fusca*. In *C. bombycina*, although several *hsp* genes were upregulated (Supplementary file S2), protein folding GO terms were not significantly enriched using our adjusted *p*-value cut-off at < 0.01 (“Protein refolding”, *p*-value= 0.025). Between *Myr. sabuleti* and *F. fusca*, the two species with a much greater number of enriched GO terms (Table S4, Fig S3), biological processes involved in mitochondria homeostasis, actin or microtubules organisation, lipid metabolism, cellular signalling and transduction, and DNA/RNA metabolism overlapped. When we categorized the differentially expressed genes into the most frequent functional categories (Fig S4), instead of only focusing on significantly enriched GO terms, we see that for all species most of the characterized differentially expressed genes are commonly comprised within the same categories being the larger proportion of them involved with cytoskeletal rearrangement and DNA/RNA metabolism. Notably, *C. holgerseni* and *Mel. bagoti* did not differentially regulated any gene involved in chaperoning and autophagy. Thus, for the 4 species with a wider gene regulation response (*i.e*., the desert species *C. bombycina* and *O. robustior*, as well as the temperate species *Myr. sabuleti* and *F. fusca*), there was a conserved differential regulation involving the same genes and pathways in response to heat stress.

To better visualise the metabolic and biochemical pathways of heat-shock response, we analysed the KEGG pathways of DET (FCx2) for each species and checked the biological function of each KEGG ortholog (KO) assigned. Overall, *C. bombycina* (7 KO), *C. holgerseni* (4 KO) and *Mel. bagoti* (5 KO) displayed few KEGG pathway matches. *C. bombycina* and *C. holgerseni* exhibited an upregulation of glutathione synthesis through its precursor [for *C. bombycina:* glycine and cysteine/methionine metabolism; for *C. holgerseni:* cysteine-glycine synthesis]. In addition, *C. holgerseni* showed a match for C10-20 isoprenoids biosynthesis (M00367). In contrast, 51 KO were assigned for *O. robustior* including genes involved in glutathione synthesis or its precursors (K00799, K00830 or K00273), lipid metabolism [Inositol phosphate metabolism (M00130, M00131, M00132), fatty acid synthesis and elongation (M00082 and M00083), and triacyl glycerol metabolism (M00089)], sugar metabolism [glucose metabolism (K01189 and K00008), trehalose synthesis (K16055 and K00963), pentose phosphate pathway (K00115 and K01623)], proline synthesis (K0086) and lysine degradation (K01435). For *F. fusca* and *M. sabuleti*, we found 88 KO and 148 KO, respectively. In both species, KO included genes involved in sugar metabolism [glycolysis (M00001) and citrate cycle (M00010)], pentose phosphate pathway (M00165 and M00004), NADP/NADPH recycling (M00172), fatty acid elongation (M00083 and M00085), tryacyl glycerol and ceramide synthesis (M00089 and M00094 respectively and methionine salvage pathway (M00034). In addition, *F. fusca* showed an enrichment in acyl glycerol and leucine degradation (M00098 and M00036) and *Myr. sabuleti* in inositol phosphate metabolism and ketone synthesis (M00130 and M00088). Overall, these results highlight the relevance of energetic pathways in ants heat stress response.

### Greater gene expression changes are associated with increased participation of taxonomically restricted genes in the heat stress response

As documented above, cross-species comparisons of gene annotation and functional categories showed that conserved genes and pathways are largely involved in the heat stress response. Nevertheless, our results also revealed several uncharacterized genes that were differentially regulated suggesting that new genes may also play a role in heat adaptation (Fig S4). To further investigate this issue, we accessed the transcripts orthology based on their protein sequence (using OrthoFinder) and considered lineage specific orthogroups as taxonomically restricted, hence, new genes. From the 292,232 protein coding genes analysed across all 6 species studied, 266,366 (91.1%) were assigned to at least one of the 34,135 orthogroups identified. Species-specific orthogroups accounted for 9,178 (26.9%) of all orthogroups and contained 29,645 genes (10.1% of the total), while 8,753 orthogroups contained all species and only 321 were single copy (Table S5). Compared to genomic orthology estimations [31], the transcriptomic based orthology analyses resulted in more species-specific and less single copy orthogroups, which can be expected since transcriptomic data include multiple isoforms per gene (see BUSCO duplication ratio at Table S2). For this reason, we estimated the relevance of new genes not only based on their frequency but comparatively, by contrasting the relative proportion of transcripts in each orthogroup category between the transcriptome and among the DET. We expected that any bias in orthology estimation due to the transcriptomic nature of the data affects the transcriptome and its subsets in a similar manner. Thus, our null hypothesis was that the proportion of transcripts in each orthology category would be equivalent in the entire transcriptome and among the differentially expressed transcripts.

According to the OrthoFinder resulting table (Supplementary file S21), we filtered orthogroups from five categories: I- orthogroups shared by all ants, II-orthogroups shared only by Myrmicinae ants, III-orthogroups shared only by Formicinae ants, IV-orthogroups shared only by the two *Cataglyphis* species, and V- species-specific orthogroups. Categories I to III included 8,753, 1,055 and 324 orthogroups respectively, and were considered as representative of taxonomically conserved genes. While categories IV and V represented taxonomically restricted genes. Category IV included 890 orthogroups. In category V, the number of species-specific orthogroups found for each species varied from 712 in *C. bombycina* to 3,053 in *Myr. sabuleti* (all species values are given in Table S5). In species with medium and high gene regulation responses, namely *C. bombycina, O. robustior, Myr. sabuleti* and *F. fusca*, we found an overall increase in the proportion of taxonomically restricted genes amongst the DET (Fig 6). This increase was significant in *F. fusca* (Fisher’s Exact Test *p*-value < 2.2e-16), *Myr. sabuleti* (*p*-value < 2.2e-16) and *O. robustior* (*p*-value < 4.5e-06), but not in *C. bombycina*, neither in *Cataglyphis* specific nor species-specific orthologs (*p*-values = 0.732 and 0.086, respectively). Contrastingly, a reduction of taxonomically restricted genes among the differentially expressed transcripts was observed in *C. holgerseni*, and in *Mel. Bagoti* only a slight increase of less than 1% in the ratio of new genes among the differentially regulated genes was found (Fig 6).

**Fig 6.**
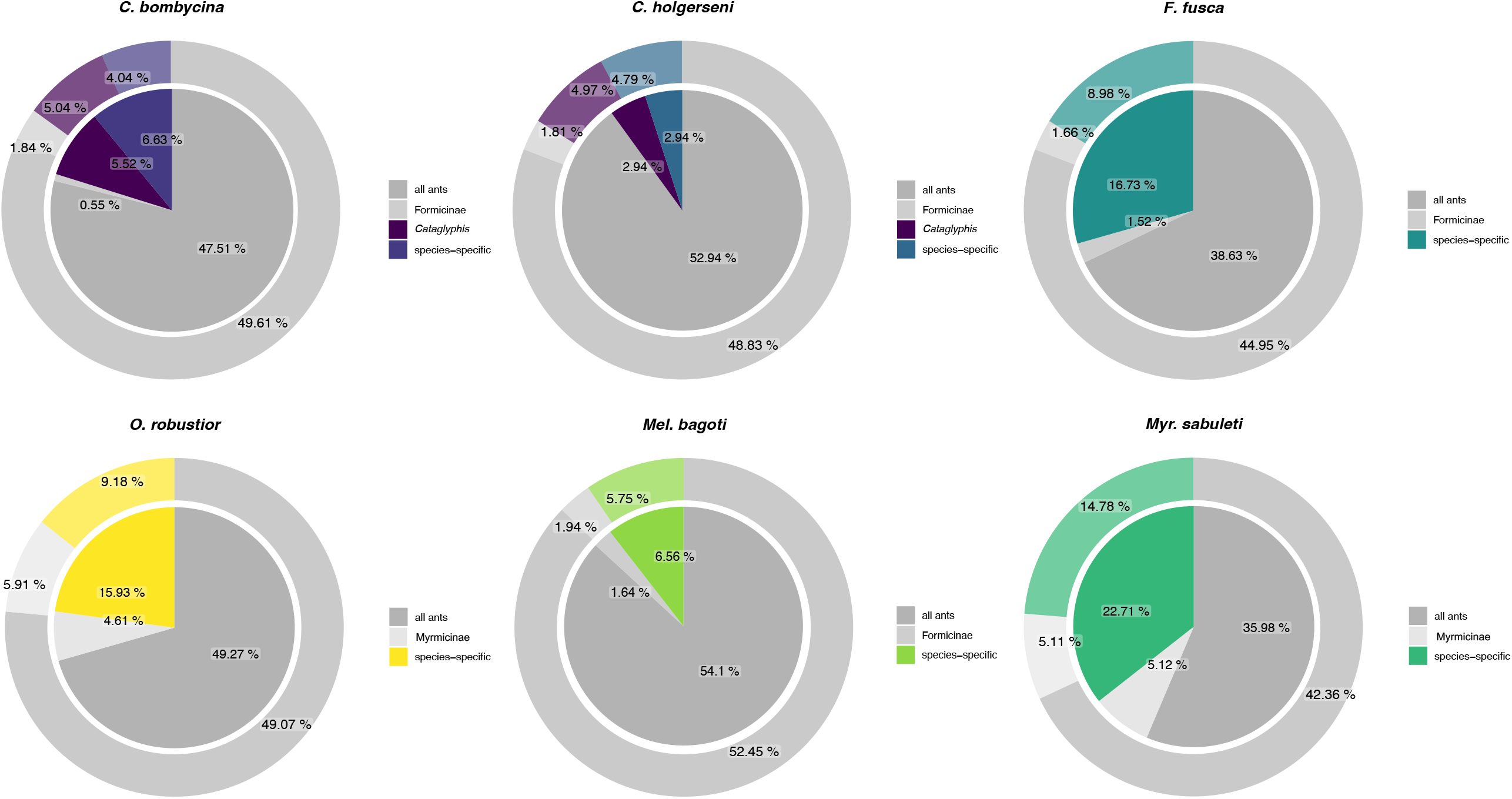
Relative proportion of transcripts in different orthologous categories. Colours highlight lineage specific categories while grey shades illustrate taxonomically conserved orthologous categories. Inner circles represent the proportion of transcripts in each category among the set of differentially expressed transcripts while outer circles show these proportions in the full transcriptome for comparison. Differences between the proportions observed in *F. fusca, Myr. sabuleti* and *O. robustior* are significant based on alpha level of .01.

These findings show that the involvement of taxonomically restricted genes in ants heat stress response is associated with the number of genes under differential expression, with species that differentially regulate more genes as temperature increases also altering the expression of a higher proportion of taxonomically restricted genes.

## Discussion

We characterized and compared the heat tolerance and the heat stress transcriptomic response of four ecological equivalent desert ant species from three distinct genera, and of two outgroup species living in a temperate region. From all 6 studied species, the red honey ant *Melophorus bagoti* was the most heat tolerant, withstanding the highest temperatures ever reported in ants [10,32–34]. However, in natural habitats, the Sahara silver ant *Cataglyphis bombycina* is the species known to endure the most environmental heat, with a mean temperature in the warmest quarter of 30.8°C and maximum temperature of 42.3°C (Table S1). An important part of *C. bombycina* heat resistance mechanism in their natural habitats relies on prism-shaped hairs covering the body of workers. These hairs reflect solar radiation through total internal reflection, which limits heat absorption when the workers are exposed to sunlight [35,36]. This adaptive mechanism deflects the energy associated with visible wavelengths but played no role in laboratory conditions when we measured heat-tolerance through infrared warming, which could explain *C. bombycina* reduced thermal tolerance in our tests (see also [10]. Not surprisingly, the least heat tolerant species studied were those from temperate areas, namely *Myrmica sabuleti* and the dusky ant *Formica fusca*. Both species displayed the lowest upper thermal limit and LT50_3h values (respectively, UTL_3h: 39°C and 41°C, LT50_3h: 39.9°C and 42°C). In ants and other insects, there is a clear association among conditions at their microclimatic level, operative temperatures experienced by individuals, and upper-thermal limits [37–39]. For example, species occupying niches exposed to greater variability in environmental temperatures, such as open areas and canopies, often show increased heat tolerance [40–42]. Our results show this association seems to also occur in strict desert thermal scavenging ants, species whose peculiar thermal niche has likely driven to the evolution of higher upper thermal limits. These observations highlight the vulnerability of ectotherms, including social insects, to temperatures changes (reviewed in [43], [2,44]).

Temperate species differentially regulated thousands of transcripts more than thermophilic ones, showing that focussed gene expression under heat stress is an important adaptation of desert ants. A similar adaptative response has also been documented at the populational level; studies comparing same species populations from colder and hotter environments found that heat adapted populations show a reduced gene expression response under heat stress. Japanese mantis shrimp populations from temperate regions differentially regulate almost one thousand more transcripts in response to heat stress than shrimp populations inhabiting a warmer environment [45]. Similarly, populations of the Chinese minnow fish from cooler regions in the north show more temperature responding genes under heat stress than southern populations [46]. In the oilseed rape (*Brassica napus*), the heat-sensitive genotype showed larger variation in the methylation pattern of multiple genes under heat stress in comparison to the heat-tolerant genotype [47]. Since molecular damage induce the heat shock response [7], reduced gene expression in heat adapted organisms might simply reflect a limited molecular damage at higher temperatures. Selection is expected to favour proteins optimized to certain thermal environments [48–50], thus the proteome of thermophile desert species might keep protein function even at elevated temperatures delaying the molecular heat shock response. A recent study on *Mytilus* mussels indeed showed that small differences in habitats were sufficient to cause such protein structural alterations [51]. Additionally, we found in desert ants many differentially expressed genes in response to heat stress that were related to DNA/ RNA metabolism and regulation (Fig S4). This is an indication that protective mechanisms against uncontrolled gene expression, such as DNA heterochromatinization and histone aggregation, are also in place. Altogether, these results suggest that the reduced gene expression alterations observed in desert ants in comparison to temperate species likely stem from a combination of diminished molecular damage and increased gene expression control mechanisms at higher temperatures, rather than from the activation of a single gene or pathway that evolved exclusively in desert ants. In line with this, the transcriptomic response of *F. fusca* and *Myr. sabuleti* involved many of the same cellular pathways and protective mechanisms found differentially regulated in desert species. Thus, temperate species have most of the molecular toolkit necessary for the heat stress response of desert ants. However, unlike the latter, temperate species co-activate many other genes and pathways upon heat stress likely leading to an unspecific and energetically costly response.

Our results suggest that desert ants evolved two adaptive strategies at the transcriptomic level to endure heat stress: (A) differentially regulating very few genes in response to heat stress, as shown by *C. holgerseni* and *Mel. bagoti*, or (B) differentially regulating several genes upon increased temperature, as observed in *C. bombycina* and *O. robustior*. In our analyses, these desert species that showed the stronger expression response under heat stress also displayed larger variation in temperature survival, with their survival curve gradually decreasing from 100 to 0% within a six-degree interval (Fig 2). Contrastingly, survival of species with fewer gene expression differences between heat stressed and non-heat stressed conditions decreased more abruptly reaching 0% within a two-degree variation interval. In face on these results, our hypothesis is that desert species with larger expression differences have a reactive response to heat, while desert species with fewer changes in gene expression keep a constitutive response to heat stress. Agreeingly, HSP70 and HSP90 transcripts are highly expressed in non-heat stressed as well as in heat stressed workers of *C. holgerseni* and *Mel. bagoti* (Fig 5). This constitutive synthesis of thermoprotective molecules would provide high heat resistance, but low flexibility at their thermal limit capacity since gene expression would not be as responsive to environmental temperature changes. Interestingly, the constitutive and reactive responses were not phylogenetically conserved (Fig 4), instead they evolved convergently in different desert species genera. Similarly, in some yeast species the expression of stress-related genes is induced upon stress while in other species these genes evolved to be constitutively activated independent of the stressor exposure [52]. Other *Cataglyphis* species studied previously also show a reactive response, similarly to *C. bombycina* and *O. robustior*, with a maximum of 1,118 DET (mean: 561 DET) accompanied by a gradual decrease in survival upon a larger temperature range [10].

Both strategies – the reactive response and the constitutive response – result in an incredible heat resistance in desert ants and could have evolved from distinct evolutionary constraints. Examples of evolutionary constraints that could lead to such differences include micro habitat differences, body-size, and metabolic trade-offs. Most *C. holgerseni* and *Mel. bagoti* foragers are larger, while *C. bombycina* and *O. robustior* workers are comparatively smaller (Fig S5). Large body sizes in desert ants could be advantageous to maintain high thermal inertia, to store more water preventing desiccation, and to store energetic resources to face cellular stress [32,53,54]. It is therefore possible that for ant species with smaller body size, a reactive gene expression response would be necessary due to reduced internal storage for keeping thermoprotective molecules. It is also important to consider that these two strategies likely involve different energetic demands. The maintenance of constitutive levels of thermoprotective genes is energetically costly, however a reactive gene expression response as temperature increases would require high amounts of energy available to be used at once to rapidly activate gene expression and to recover from the possible loss of biochemical activity from protein denaturation [55]. Therefore, energetic trade-offs and alterations in energetic pathways could represent important constraints to the evolution of these two strategies, as is suggested by the activation of energetic pathways in the heat stress response.

Amongst desert ants, in *C. holgerseni* and *M. bagoti* we found respectively 3 and 4 differentially expressed transcripts involved in sugar energetic pathways. Interestingly, in *O. robustior* more than 20 transcripts were upregulated in this functional category. For example, we found genes involved in glucose synthesis (*maltase 1-like, alpha-glucosidase* or *alpha-amylase a-like*) and transformation (*l glucose dehydrogenase fad quinone-like* or *sorbitol dehydrogenase*). In addition, KEGG pathway analyses revealed that among the DET of *O. robustior* there was a positive regulation of trehalose synthesis and of the pentose phosphate pathway (PPP). The former is known to enhance heat and desiccation tolerance in various organisms [56], while the oxidative phase of PPP is used to regenerate NADPH which is required by antioxidant enzymes such as the *glutathion S-transferase* and the *cytochrome P450*, both also upregulated in *O. robustior* [57,58]. A positive co-regulation of sugar metabolism in response to heat stress was previously documented in *Cataglyphis* [10]. Similarly, among the temperate species, many transcripts differentially expressed are involved in sugar metabolism as well as the upregulation of the PPP pathway. These results suggest that the heat-shock response in ants requires a lot of energy provided by the sugar metabolism. Nevertheless, in *O. robustior* this response was somewhat accentuated, with many of the genes and regulatory pathways involved in the antioxidative machinery along with genes involved in the trehalose synthesis being differentially regulated under heat stress.

The overlap of genes and pathways involved in the transcriptomic response to heat stress of desert and temperate ants is strong evidence of taxonomically conserved genes and pathways regulating this trait. Nevertheless, our analysis also shows an increased proportion of new genes amongst the DET of desert and non-desert species when compared to the proportion of new genes in their entire transcriptome (Fig 6). Only desert ants with fewer gene expression alterations were an exception to this pattern. These observations indicate that new genes are likely to be co-expressed along with taxonomically conserved genes under heat-stress, possibly because they share the same promoter regions [59]. Agreeingly, in our data, the larger the number of genes under differential expression, the larger the proportion of new genes amongst them. This co-expression offers an opportunity to gene neofunctionalization during adaptation, which is one of the most important origins of new genes fixation [60,61]. New genes are known to be particularly relevant to species adaptative response [61–64]. For example, in Antarctic notothenioid fishes, the ice-biding antifreeze glycoprotein (AFGP), that originated from a functionally unrelated gene (the *pancreatic trypsinogen-like protease2*), has been co-opted to play a crucial adaptive role in preventing blood from freezing [65]. In ants, 10% to 30% of all genes in the genome are estimated to be taxonomically restricted [66], and evidence suggest that high ratios of *de novo* gene formation exist within the group [67]. It is therefore possible that taxonomically restricted genes are also involved in species-specific adaptations to heat stress in desert ants. Comparative genomic studies and functional analyses should further explore these genes and their putative roles in heat stress adaptation.

Interspecies comparative transcriptomic analyses are challenging, because normalization methods cannot account properly for species specific batch effects and the use of phylogenetic information to weight transcriptomic differences can be misleading [68]. Herein, we first performed an intra species transcriptomic analysis and then compared the results across species in search of similarities/differences. This approach has the advantage of avoiding normalization problems across species – particularly when using whole body data –, but it has the caveat of accumulating trade-off errors of the independent analyses, as for each species test, we would expect independent false positive and false negative results. Due to our conservative data analyses pipeline (overlapping two programs for differential analyses, using conserved FDR and FC cut-offs), we expect our results to be mostly impacted by false negatives. Thus, transcripts differentially regulated between the two conditions may not be detected as so statistically, and consequently, matches of common DET across species will be reduced. Moreover, we have chosen to pool whole body samples from multiple individuals and colonies to reduce individual variation biases and to focus on overall changes in expression tendencies. That means that transcripts with reduced expression and/or with tissue specific variations were overseen in our results. We also found that the quality of the transcriptome assembly was affected by using the reference genome, thus species whose genomes were available had less fragmented transcriptome assemblies, as suggested by the quality analyses in Table S2. This could affect our results in two ways, first by reducing the gene counts of fragmented transcripts in certain species, thus hindering their identification as differentially expressed, and secondly by interfering with transcripts annotation. Based on the differences of BUSCO completeness between transcriptomes with and without the genome guided assembly, we expect that around 10% of the transcripts could be affected in this way. Still, we have chosen to use the reference genomes to improve some assemblies, first because we had at least one desert and one temperate species with a reference genome, allowing us to account at some level for the impact of different assembly strategies, and secondly because we wanted to make available to the community the best reference transcriptome sets possible. These caveats could hardly be avoided due to the before mentioned reasons, and overall, they do not hinder the main conclusions of our study.

In summary, our results show that, as opposed to temperate ant species, desert ants have reduced gene expression alterations even when under extreme heat stress. Furthermore, they support the hypothesis that desert species evolved convergently two transcriptomic strategies to withstand heat stress: in *C. holgerseni* and *Mel. Bagoti*, a constitutive level of thermoprotective genes reduces the need for great changes in gene expression when temperature increases; contrastingly, in *C. bombycina* and *O. robustior* a larger number of genes are differentially regulated in response to increasing heat. These two molecular responses are associated with the survival pattern of the ants: the species with fewer gene expression alterations have a small variability in individual survival, with all workers dying within a two-degree interval, while species with greater expression responses experience a survival reduction from 100% to 0% within a larger temperature interval of six degrees. The interplay among body size, energetic metabolism, and heat tolerance in the evolution of these two strategies deserves further exploration. Overall, the heat stress response involved mostly conserved genes and pathways across the studied species, still new genes were co-expressed during heat stress offering an opportunity for adaptative gene neofunctionalization. Finally, this study shows that, in some circumstances, transcriptional differences between conditions may be insufficient to understand putative adaptative mechanisms if they are not contextualized within a phylogenetic perspective. Indeed, the expression of some genes may be constitutive and, therefore, these genes will not be detected as differentially regulated even though they may be functionally relevant to species adaptation. Clearly, transcriptome studies should consider an evolutionary approach for comparative data analyses including multiple species from different phylogenetical backgrounds.

## Material and methods

### Ant sampling and rearing conditions

We studied six ant species from different locations. Four from arid and sandy deserts: *Cataglyphis bombycina* (Morocco), *C. holgerseni* (Israel), *Ocymyrmex robustior* (Namibia) and *Melophorus bagoti* (Australia); and two from a temperate region: *Formica fusca* and *Myrmica sabuleti* (Belgium). The four desert species can be considered ecologically equivalent [43]. They inhabit sandy desert environments with mean annual temperature (AMT) above 21°C, and warmest month recorded (MaxT) ranging from 32.3 to 42.3 °C (Table S1). We included two species of desert ants from the same genus (*C. bombycina* and *C. holgerseni*) to account for intra genus variation, as well as species from three different genera of heat adapted ants (*Cataglyphis*, *Ocymyrmex* and *Melophorus*) that evolved their adaptations independently to account for inter genera variability. The two temperate ant species, *Formica fusca* and *Myrmica sabuleti*, were chosen as comparative outgroups being, respectively, representative species from the Formicinae and the Myrmicinae subfamilies. Both these species inhabit temperate areas with AMT of 10.2°C and MaxT of 22.8°C. Colonies of desert species are headed by a single queen (monogyny); queens are polyandrous (multiple mated), except for *O. robustior* where queens are strictly monandrous [69–72]. The two temperate species studied are facultatively polygynous (a single or several reproductive queens per colony); *F. fusca* is facultatively polyandrous, no information about queen mating frequency is available for *Myr. sabuleti* [73,74].

We excavated 4–6 colonies for each species at sampling sites and brought them back to the lab accompanied by the queen, brood, and workers. The colonies were kept under steady conditions: mean temperature of 25°C (± 1°C) for desert species and 18°C (± 1°C) for temperate species, light-dark cycle of 12 h/12 h, and relative humidity between 30% and 40% for all ants. Environmental conditions during this adaptation period were established based on species optimal parameters for laboratory brood rearing (personal observations). Ants were fed with sugar solution provided *ad libitum*, and sliced mealworms that were provided three times per week. Colonies were maintained under these conditions for one month prior to the beginning of the assays. Only adult workers with a highly sclerotized and melanized exoskeleton (*i.e*., likely born in the field) were used.

### Heat tolerance assays

Workers heat tolerance was determined using a heat stress assay, as detailed in [10,22]. For each species, six groups (*n* = 6) of 10 randomly selected workers (from 4-6 colonies per species) were placed in glass tubes – totalizing 60 workers – and used in the heat tolerance assays. Because body size influences heat tolerance due to differences in relative water loss [75–77], a wet cotton ball was added in the testing tubes to reduce ant desiccation and its effect on specimens’ survival during the tests. The tubes were immersed in a water bath (SW22, Julabo GmbH, Seelbach, Germany) kept at constant temperature of 39°C, 41°C, 43°C, 45°C, 47°C, 49°C or 51°C. Tubes temperatures were monitored using 0.075-mm diameter thermocouples (Type K Thermocouple [Chromel/Alumel], RS Components Ltd, UK) connected to a digital thermometer (RS Pro RS52 Digital Thermometer, RS Components Ltd, UK). Percent survival was recorded after 3 hours; workers were identified as moribund once they lost their locomotor ability (*i.e*., muscular paralysis; [78]). For each species, we measured two indicative parameters of heat tolerance: (A) the upper thermal limit (UTL_3h), the temperature below which the survival rate is significantly different from 100% (ANOVA test followed by a Tukey’s test; *p* < 0.05), and (B) the median lethal temperature (LT50_3h), the temperature at which survival probability equals 50%. LT50_3h was estimated using a simple logistic regression of death probability in function of the temperature. The LT50_3h, and its 95% confidence intervals, were compared between species using the ratio test [79], implemented in the *ratio_test* function from the *ecotox* package [80].

### RNA-Seq library preparation and sequencing

For each species, we sequenced the total RNA of four replicates per treatment (*n* = 4). In the heat stress (HS) treatment, workers were exposed to their UTL_3h for 3 hours; in the non-heat stress (NHS or control) treatment, workers were maintained in the glass tubes for 3 hours at 25°C or 18°C, for desert and temperate species respectively. Each sample replicate contained 10 workers randomly chosen among the different colonies of a species (4-6 colonies per species), totalizing 40 workers per treatment among 4 replicates. Ants were immediately frozen after treatments and stored at −80°C until RNA extraction. Workers were pooled to avoid individual and colony biases expression responses. Since heat stress may affect multiple body tissues, we used entire bodies for expression analyses. RNA extractions, RNA quality assessments (determined using Bioanalyzer), RNA-seq library preparations and sequencing were performed by BGI Tech Solutions (Hong Kong). Total RNA was extracted using Trizol (Invitrogen, Carlsbad, CA, USA) according to manufacturer’s instructions (EDTA 250mM was added to *Formica fusca* extractions to avoid non enzymatic RNA degradation; Valles et al., 2012). Sequencing was performed using an Illumina HiSeq 4000 System. About 25 million single reads of 50 bp in length were generated per sample replicate. We additionally sequenced for each species a pool of workers (HS and NHS), eggs, and pupae using a HiSeq X Ten System that was used to assemble the species reference transcriptomes. About 90 million paired reads of 150 bp in length were obtained for each of these species’ pools.

### Transcriptome assembly and differential expression

Transcriptome assembly and differential expression analyses were performed independently for each species. For both *Cataglyphis* species, the transcriptome and differential expression data have been already published elsewhere using the same methodology described hereafter [10]. Quality of all sequenced reads were estimated using FastQC v0.11.7 [81]. We used the Trinity v2.8.4 pipelines [82] with the sequenced dataset from the pool of workers, eggs and pupae for transcriptome assemblies. To increase assembly efficiency, reads were first digitally normalized (20x coverage) through computational coverage systematization [83]. Then two assembly strategies were used depending on the species: a full *de novo* and a reference guided assembly. For the two *Cataglyphis* species and for *Formica fusca* we generated both, the full *de novo* and the reference guided assemblies. The reference guided approach for the *Cataglyphis* was based on the genome of *Cataglyphis hispanica* (assembly ASM419527v1 - [31] and that of *F. fusca* in the genome of *Formica exsecta* (assembly ASM365146v1 - [84]. These two independent assemblies were then concatenated. For *Ocymyrmex robustior*, *Melophorus bagoti* and *Myrmica sabuleti* only the full *de novo* assembly was performed due to lack of a close reference genome. In all cases, transcripts redundancy was initially removed with CD-Hit v4.8.1 [28] using the threshold of 95% nucleotide similarity, then Corset v1.08 [29] and Lace v1.13 [30] were used to cluster transcripts into superTranscripts, which further reduced transcript redundancy and improved posterior gene expression counts [30]. Assemblies were then annotated with Annocript v2.0.1 [85] in tandem with the UniProt Reference Clusters (UniRef90) and the UniProtKB/Swiss-Prot [86] databases (March 2019 versions). Only transcripts potentially encoding proteins – based on ORF estimation or blast results – were retained in the final reference transcriptome assemblies. All the parameters used in the assembly and annotation pipelines were the programmes’ suggested defaults, unless otherwise stated. Assessments of assemblies’ quality were obtained with BUSCO v5.3.2 [87], against the Hymenoptera (odb10) database, and with QUAST v5.0.2 [88].

For the differential gene expression analyses, reads from ants of the HS and NHS treatments were aligned to the reference transcriptome and counted using Salmon [89]. The transcripts that were differentially expressed between treatments were identified using the edgeR [90] and the DESeq2 [91] pipelines. To increase stringency and reduce the rate of false positive results, the differentially expressed transcripts were selected based on the overlapping results of both analyses (*i.e*., overlapping results from edgeR and DESeq2) and using a cut-off of mean absolute log_2_-fold change ≥ 2 between treatments and FDR ≤1e-3 (as suggested in the Trinity pipeline for *de novo* transcriptomes). We then computationally compared the list of differentially expressed transcripts between each species pair based on their UniRef90 gene name annotation using R (*stringr* library). Only unique, non-redundant and meaningful gene annotations (*i.e*., genes whose annotation did not contain “uncharacterised protein”) were included in this comparison and in the statistical tests. After this computational comparison of gene names, the final list of overlapping genes was manually curated to avoid partial or redundant annotation matches. Significance of common differentially expressed genes between species pairs was assessed using 10,000 random samplings comparisons based on the transcriptome datasets, i.e., the same number of DET was randomly sampled from the transcriptome of each species and compared with another species randomly sampled in the same manner. These comparisons were repeated 10,000 times per species pair and a cut-off *p*-value < 0.01 of finding the same number of common DET by chance was used (Rscript available at https://github.com/nat2bee/Foragers_vs_Nurses/blob/master/Statistics/common.stats.R.

### Functional analyses of the differentially expressed genes

To test whether gene ontology (GO) terms for biological processes (BP) were enriched among the differentially expressed transcripts, we kept only transcripts displaying non-zero expression in the treatment groups (i.e., for *C. bombycina:* 29,965 transcripts; for *C. holgerseni:*26,769 transcripts; for *Mel. bagoti:* 38,726 transcripts; for *O. robustior:* 45,701 transcripts; for *M. sabuleti:* 84,784 transcripts; and for *F. fusca:* 62,416 transcripts). We used the “weight01” algorithm with the Fisher enrichment test in the R package TopGO [92] for the enrichment test, and significance was evaluated based on an adjusted alpha level of 0.01 to obtain only highly significant enriched terms. For summary visualization of enriched GO terms in desert and non-desert species, we used the Revigo server and the *Drosophila* database [93].

For each species, the function of all differentially expressed transcripts was manually checked and categorized according to their metabolic and cellular role from which we selected nine most frequent categories: Apoptose, Authophagy, Chaperoning, Cytoskeletal rearrangement, DNA/RNA metabolism, Lipid metabolism, Proteolysis, ROS elimination, Sugar metabolism, Other and Uncharacterized (*i.e*., genes of unknown function or not annotated). Finally, we assessed gene involvement in specific biochemical and metabolic pathways using the Kyoto Encyclopedia of Genes and Genomes (KEGG), via the KEGG Automatic Annotation Server (KAAS; [94]).

### Expression levels of Heat shock proteins

We filtered transcripts in the reference transcriptome annotated to the heat shock protein genes HSP70 and HSP90 using a custom script (Find_Genes.py available at https://github.com/nat2bee/Cataglyphis_HS/tree/main/Scripts) based on representative search terms annotation (*i.e., Hsp70, Hsp 70, Heat shock protein 70, Hsp90, Hsp 90, Heat shock protein 90*). For each of these transcripts, we then plotted their normalized expression count in TMM from all heat stressed and non-heat stressed replicates for their comparative visualization.

### Orthologous and new genes identification

We used TransDecoder v5.5.0 [95] to convert transcriptome ORF regions to amino acid sequences. Using OrthoFinder v2.5.4 [96] with default parameters, we then estimated orthology among the transcriptomes of all species and generated the species phylogenetic tree using MAFFT for sequence alignment and FasTree for tree estimation. Orthogroups containing at least one transcript differentially expressed were considered as a differentially expressed orthogroup. We classified the orthogroups into five categories filtering the OrthoFinder resulting table: I-orthogroups containing representative transcripts from all ants, II-orthogroups containing representative transcripts only in Myrmicinae ants, III-orthogroups containing representative transcripts only in Formicinae ants, IV-orthogroups containing representative transcripts only in the two *Cataglyphis* species, accounting for intra genus similarities and V-species-specific orthogroups. Transcripts within orthogroups in categories IV and V were considered as taxonomically restricted genes, while categories I, II and III were composed by conserved orthologous. For each species, only the relevant categories were considered. Fisher’s Exact Test was used to test whether the proportion of genes in each of these categories diverged significantly between the differentially expressed transcripts and the entire species transcriptome.

## Data availability

Sequenced data and reference transcriptome assemblies for the studied ant species are available at NCBI under the Bioproject PRJNA632584; the annotations for these assemblies and scripts used are accessible at GitHub (https://github.com/nat2bee/Cataglyphis_HS).

## Acknowledgments

This work was supported by the Belgian *Fonds National pour la Recherche Scientifique* (grant numbers J.0151.16 to SA, 40005980 to NSA and T.0140.18 to RP) and the Universté Libre de Bruxelles (FER 2017 and 2021). We are grateful to Morocco’s Département des Eaux et Forêts (permit number: HCEFLCD - Decisions 11/2018, 23/2019 and 13/2020 *DEF/DLCDPN/DPRN/CFF*), to the Namibian National Commission on Research, Science and Technology (permit number: RPIV01042022), to the Australian Department of Environment and Energy (permit number: PWS2018-AU-002350), to the Parks & Wildlife Commission of Northern Territory (permit number: 63296) and to the Center for Appropriate Technology of Alice Springs for granting us collection and exportation permits. We thank A. Kuhn, and S. Deeti for their help with field collections.

