## Supplementary File 1 for "Facing lethal temperatures: heat shock response in desert and temperate ants"

Natalia de Souza Araujo

Department of Evolutionary Biology & Ecology, CP 160/12

Avenue F.D. Roosevelt, 50

B - 1050 Brussels

Belgium

**Table S1.** Main thermal characteristics of species natural habitats. All values are in °C. The climatic data were obtained from the WorldClim database using a resolution of 30 arc-seconds.

| Species | Annual mean temperature | Mean diurnal range | Mean temperature of warmest quarter | Maximum temperature | Coordinates |
|---|---|---|---|---|---|
| *C. bombycina* (Morocco) | 21.0 | 17.2 | 30.8 | 42.3 | 30°33′22″ N<br>-5°83′83″ E |
| *C. holgerseni* (Israël) | 22.9 | 13.1 | 29.6 | 37.3 | 30°41'22'' N<br>35°14'14'' E |
| *M. bagoti* (Australia) | 21 | 16.3 | 28.2 | 37.3 | -23°50'35'' N<br>133°57'18'' E |
| *O. robustior* (Namibia) | 21.1 | 16.3 | 23.6 | 32.3 | -23°33'45'' N<br>15°2'31'' E |
| *M. sabuleti / F. fusca* (Belgium) | 10.2 | 8.2 | 17.1 | 22.8 | 50°38'41'' N<br>4°14'22'' E |

**Table S2**. Major summary quality parameters of the reference transcriptome assemblies. BUSCOs parameters are based on a N of 5,991: C = complete; S = single copy; D = duplicated; F = fragmented; M = missing.

| | ***C. bombycina*** | ***C. holgerseni*** | ***Mel. bagoti*** | ***O. robustior*** | ***F. fusca*** | ***Myr. sabuleti*** |
|---|---|---|---|---|---|---|
| **Number of transcripts** | 41,912 | 44,525 | 38,726 | 45,701 | 62,416 | 84,784 |
| **Total length (pb)** | 115,117,532 | 112,126,054 | 85,836,189 | 91,769,795 | 146,813,706 | 139,366,828 |
| **N50** | 4,446 | 4,153 | 3,929 | 3,738 | 4,507 | 3,123 |
| **BUSCOs** | C:93.5% [S:26.9%, D:66.6%], F:2.7%, M:3.8% | C:92.9% [S:28.2%, D:64.7%], F:2.8%, M:4.3% | C:85.9% [S:34.2%, D:51.7%], F:5.0%, M:9.1% | C:85.3% [S:25.7%, D:59.6%], F:5.6%, M:9.1% | C:95.9% [S:29.4%, D:66.5%], F:1.2%, M:2.9% | C:86.7% [S:23.6%, D:63.1%], F:5.4%, M:7.9% |
| **Blast hits to Swiss-Prot** | 24,805 | 25,225 | 21,369 | 27,151 | 30,795 | 30,858 |
| **Blast hits to UniRef90** | 35,155 | 36,537 | 33,296 | 40,188 | 48,761 | 61,977 |
| **Transcripts with at least one blast hit** | 35,416 | 36,951 | 33,329 | 40,229 | 48,804 | 62,037 |

**Table S3.** Number of differentially expressed genes under heat stress common among all ant species. The number of differentially expressed genes shared by each species pair is shown in the upper right-hand part of the table. Shown in the lower left-hand part of the table are the mean numbers and the standard deviation (in parentheses) of the differentially expressed genes expected to be shared by each species pair, based on the distributions of 10,000 random iterations. The cells in grey show the numbers of unique non-redundant genes that were differentially expressed in each species. * = significant number of genes in common (*p* < 0.01).

| | ***C. bombycina*** | ***C. holgerseni*** | ***F. fusca*** | ***O. robustior*** | ***Mel. bagoti*** | ***Myr. sabuleti*** |
|---|---|---|---|---|---|---|
| ***C. bombycina*** | 113 | 2 | 42* | 15* | 2 | 40* |
| ***C. holgerseni*** | 0.83 (0.90) | 40 | 13 | 5 | 1 | 12 |
| ***F. fusca*** | 23.5 (4.52) | 8.7 (2.78) | 1,394 | 95* | 12 | 419* |
| ***O. robustior*** | 5.1 (2.20) | 1.88 (1.35) | 53.42 (6.85) | 315 | 5 | 116* |
| ***Mel. bagoti*** | 0.62 (0.79) | 0.23 (0.47) | 6.64 (2.42) | 1.44 (1.18) | 31 | 11 |
| ***Myr. sabuleti*** | 27.36 (4.76) | 10.13 (3.01) | 289.72 (14.81) | 76.65 (7.96) | 7.91 (2.63) | 2,048 |

**Table S4.** GO terms enriched among the differentially expressed transcripts under heat stress in each species.

| Species | GO ID | Term | Annotated | Significant | Expected | Classic Fisher p-value |
|---|---|---|---|---|---|---|
| *C. bombycina* | GO:1903699 | tarsal gland development | 1 | 1 | 0 | 0.0044 |
| | GO:0046697 | decidualization | 1 | 1 | 0 | 0.0044 |
| | GO:1903709 | uterine gland development | 1 | 1 | 0 | 0.0044 |
| | GO:0002674 | negative regulation of acute inflammatory response | 1 | 1 | 0 | 0.0044 |
| | GO:0050953 | sensory perception of light stimulus | 100 | 2 | 0.44 | 0.0087 |
| | GO:0007288 | sperm axoneme assembly | 2 | 1 | 0.01 | 0.0088 |
| | GO:0060426 | lung vasculature development | 2 | 1 | 0.01 | 0.0088 |
| | GO:0061038 | uterus morphogenesis | 2 | 1 | 0.01 | 0.0088 |
| | GO:0042487 | regulation of odontogenesis of dentin-containing tooth | 2 | 1 | 0.01 | 0.0088 |
| | GO:1903929 | primary palate development | 2 | 1 | 0.01 | 0.0088 |
| | GO:0001886 | endothelial cell morphogenesis | 2 | 1 | 0.01 | 0.0088 |
| | GO:0043568 | positive regulation of insulin-like growth factor receptor signaling pathway | 2 | 1 | 0.01 | 0.0088 |
| | GO:0036334 | epidermal stem cell homeostasis | 2 | 1 | 0.01 | 0.0088 |
| | GO:0034405 | response to fluid shear stress | 2 | 1 | 0.01 | 0.0088 |
| *C. holgerseni* | GO:0006879 | cellular iron ion homeostasis | 15 | 2 | 0.03 | 0.00035 |
| | GO:0006826 | iron ion transport | 18 | 2 | 0.03 | 0.00051 |
| | GO:0042773 | ATP synthesis coupled electron transport | 15 | 2 | 0.03 | 0.00535 |
| | GO:0051638 | barbed-end actin filament uncapping | 4 | 1 | 0.01 | 0.00762 |
| | GO:0050830 | defense response to Gram-positive bacterium | 4 | 1 | 0.01 | 0.00762 |

| | | | | | | |
|---|---|---|---|---|---|---|
| | GO:2000813 | negative regulation of barbed-end actin filament capping | 4 | 1 | 0.01 | 0.00762 |
| | GO:0006122 | mitochondrial electron transport, ubiquinol to cytochrome c | 5 | 1 | 0.01 | 0.00951 |
| ***Mel. bagoti*** | GO:0022904 | respiratory electron transport chain | 35 | 4 | 0.12 | 0.00046 |
| | GO:1904749 | regulation of protein localization to nucleolus | 2 | 1 | 0.01 | 0.00658 |
| ***O. robustior*** | GO:0031047 | gene silencing by RNA | 72 | 7 | 1.21 | 5.5e-06 |
| | GO:0042026 | protein refolding | 9 | 4 | 0.15 | 9.0e-06 |
| | GO:0034605 | cellular response to heat | 23 | 5 | 0.39 | 3.3e-05 |
| | GO:0006457 | protein folding | 89 | 11 | 1.5 | 0.00024 |
| | GO:0006275 | regulation of DNA replication | 20 | 4 | 0.34 | 0.00030 |
| | GO:0051085 | chaperone cofactor-dependent protein refolding | 4 | 2 | 0.07 | 0.00165 |
| | GO:0006265 | DNA topological change | 18 | 3 | 0.3 | 0.00317 |
| | GO:0071923 | negative regulation of cohesin loading | 7 | 2 | 0.12 | 0.00560 |
| | GO:0051312 | chromosome decondensation | 7 | 2 | 0.12 | 0.00560 |
| ***F. fusca*** | GO:0007169 | transmembrane receptor protein tyrosine kinase signaling pathway | 211 | 38 | 16.4 | 2.2e-12 |
| | GO:0042026 | protein refolding | 18 | 12 | 1.4 | 5.4e-10 |
| | GO:0034605 | cellular response to heat | 32 | 14 | 2.49 | 5.0e-08 |
| | GO:0006458 | 'de novo' protein folding | 14 | 11 | 1.09 | 2.7e-06 |
| | GO:0006457 | protein folding | 109 | 33 | 8.47 | 3.1e-06 |
| | GO:0051085 | chaperone cofactor-dependent protein refolding | 8 | 6 | 0.62 | 5.3e-06 |
| | GO:0045041 | protein import into mitochondrial intermembrane space | 6 | 5 | 0.47 | 1.6e-05 |
| | GO:0050808 | synapse organization | 299 | 26 | 23.24 | 7.2e-05 |
| | GO:0015074 | DNA integration | 178 | 29 | 13.84 | 0.00010 |

| | | | | | | |
|---|---|---|---|---|---|---|
| | GO:0050829 | defense response to Gram-negative bacterium | 34 | 10 | 2.64 | 0.00018 |
| | GO:1902037 | negative regulation of hematopoietic stem cell differentiation | 3 | 3 | 0.23 | 0.00047 |
| | GO:0010529 | negative regulation of transposition | 6 | 4 | 0.47 | 0.00048 |
| | GO:0008637 | apoptotic mitochondrial changes | 37 | 11 | 2.88 | 0.00050 |
| | GO:0090074 | negative regulation of protein homodimerization activity | 7 | 4 | 0.54 | 0.00105 |
| | GO:0031547 | brain-derived neurotrophic factor receptor signaling pathway | 12 | 5 | 0.93 | 0.00140 |
| | GO:2000481 | positive regulation of cAMP-dependent protein kinase activity | 12 | 5 | 0.93 | 0.00140 |
| | GO:0048022 | negative regulation of melanin biosynthetic process | 12 | 5 | 0.93 | 0.00140 |
| | GO:2001224 | positive regulation of neuron migration | 12 | 5 | 0.93 | 0.00140 |
| | GO:2001214 | positive regulation of vasculogenesis | 12 | 5 | 0.93 | 0.00140 |
| | GO:2000670 | positive regulation of dendritic cell apoptotic process | 12 | 5 | 0.93 | 0.00140 |
| | GO:0016246 | RNA interference | 22 | 7 | 1.71 | 0.00195 |
| | GO:0008050 | female courtship behavior | 8 | 4 | 0.62 | 0.00197 |
| | GO:1901888 | regulation of cell junction assembly | 153 | 17 | 11.89 | 0.00211 |
| | GO:0043950 | positive regulation of cAMP-mediated signaling | 13 | 5 | 1.01 | 0.00213 |
| | GO:0021884 | forebrain neuron development | 26 | 6 | 2.02 | 0.00310 |
| | GO:0071880 | adenylate cyclase-activating adrenergic receptor signaling pathway | 14 | 5 | 1.09 | 0.00310 |
| | GO:0030950 | establishment or maintenance of actin cytoskeleton polarity | 9 | 4 | 0.7 | 0.00332 |
| | GO:0051146 | striated muscle cell differentiation | 166 | 11 | 12.9 | 0.00341 |

| | | | | | | |
|---|---|---|---|---|---|---|
| | GO:0045089 | positive regulation of innate immune response | 46 | 12 | 3.58 | 0.00370 |
| | GO:0050774 | negative regulation of dendrite morphogenesis | 27 | 7 | 2.1 | 0.00373 |
| | GO:0046475 | glycerophospholipid catabolic process | 5 | 3 | 0.39 | 0.00415 |
| | GO:0032486 | Rap protein signal transduction | 15 | 5 | 1.17 | 0.00435 |
| | GO:0032092 | positive regulation of protein binding | 28 | 7 | 2.18 | 0.00465 |
| | GO:0048036 | central complex development | 10 | 4 | 0.78 | 0.00520 |
| | GO:0007257 | activation of JUN kinase activity | 16 | 5 | 1.24 | 0.00593 |
| | GO:0038180 | nerve growth factor signaling pathway | 16 | 5 | 1.24 | 0.00593 |
| | GO:0070868 | obsolete heterochromatin organization involved in chromatin silencing | 16 | 5 | 1.24 | 0.00593 |
| | GO:0016042 | lipid catabolic process | 134 | 16 | 10.42 | 0.00596 |
| | GO:0006336 | DNA replication-independent chromatin assembly | 11 | 4 | 0.86 | 0.00768 |
| | GO:0097345 | mitochondrial outer membrane permeabilization | 10 | 4 | 0.78 | 0.00781 |
| | GO:0042247 | estestablishment of planar polarity of follicular epithelium | 6 | 3 | 0.47 | 0.00783 |
| | GO:0015771 | trehalose transport | 6 | 3 | 0.47 | 0.00783 |
| | GO:1902463 | protein localization to cell leading edge | 6 | 3 | 0.47 | 0.00783 |
| | GO:0030033 | microvillus assembly | 17 | 5 | 1.32 | 0.00788 |
| | GO:0044331 | cell-cell adhesion mediated by cadherin | 31 | 7 | 2.41 | 0.00842 |
| | GO:0051491 | positive regulation of filopodium assembly | 47 | 9 | 3.65 | 0.00923 |
| ***Myr. sabuleti*** | GO:0042026 | protein refolding | 18 | 11 | 2.08 | 6.9e-07 |
| | GO:0006458 | 'de novo' protein folding | 22 | 11 | 2.54 | 3.7e-06 |
| | GO:1990481 | mRNA pseudouridine synthesis | 5 | 5 | 0.58 | 2.0e-05 |
| | GO:0000495 | box H/ACA snoRNA 3'-end processing | 5 | 5 | 0.58 | 2.0e-05 |

| | | | | | | |
|---|---|---|---|---|---|---|
| | GO:0031120 | snRNA pseudouridine synthesis | 5 | 5 | 0.58 | 2.0e-05 |
| | GO:0031118 | rRNA pseudouridine synthesis | 5 | 5 | 0.58 | 2.0e-05 |
| | GO:0045041 | protein import into mitochondrial intermembrane space | 13 | 8 | 1.5 | 2.3e-05 |
| | GO:0034605 | cellular response to heat | 31 | 12 | 3.58 | 9.2e-05 |
| | GO:0008637 | apoptotic mitochondrial changes | 37 | 11 | 4.28 | 0.00011 |
| | GO:0035220 | wing disc development | 130 | 23 | 15.02 | 0.00046 |
| | GO:0006000 | fructose metabolic process | 5 | 4 | 0.58 | 0.00081 |
| | GO:0006003 | fructose 2,6-bisphosphate metabolic process | 5 | 4 | 0.58 | 0.00081 |
| | GO:0007614 | short-term memory | 29 | 10 | 3.35 | 0.00103 |
| | GO:0016567 | protein ubiquitination | 221 | 30 | 25.54 | 0.00105 |
| | GO:0007281 | germ cell development | 304 | 39 | 35.13 | 0.00117 |
| | GO:0044351 | macropinocytosis | 3 | 3 | 0.35 | 0.00154 |
| | GO:0040040 | thermosensory behavior | 13 | 6 | 1.5 | 0.00196 |
| | GO:0006610 | ribosomal protein import into nucleus | 19 | 7 | 2.2 | 0.00387 |
| | GO:0019941 | modification-dependent protein catabolic process | 341 | 43 | 39.41 | 0.00464 |
| | GO:0046084 | adenine biosynthetic process | 7 | 4 | 0.81 | 0.00465 |
| | GO:0031291 | Ran protein signal transduction | 7 | 4 | 0.81 | 0.00465 |
| | GO:0001654 | eye development | 185 | 21 | 21.38 | 0.00472 |
| | GO:0010886 | positive regulation of cholesterol storage | 4 | 3 | 0.46 | 0.00563 |
| | GO:0034383 | low-density lipoprotein particle clearance | 4 | 3 | 0.46 | 0.00563 |
| | GO:0030953 | astral microtubule organization | 4 | 3 | 0.46 | 0.00563 |
| | GO:0043097 | pyrimidine nucleoside salvage | 4 | 3 | 0.46 | 0.00563 |
| | GO:1901741 | positive regulation of myoblast fusion | 4 | 3 | 0.46 | 0.00563 |
| | GO:0061512 | protein localization to cilium | 16 | 6 | 1.85 | 0.00675 |
| | GO:0007619 | courtship behavior | 58 | 15 | 6.7 | 0.00711 |

| | | | | | | |
|---|---|---|---|---|---|---|
| | GO:0046958 | nonassociative learning | 24 | 7 | 2.77 | 0.00842 |
| | GO:0006465 | signal peptide processing | 8 | 4 | 0.92 | 0.00846 |
| | GO:0006198 | cAMP catabolic process | 8 | 4 | 0.92 | 0.00846 |
| | GO:0010738 | regulation of protein kinase A signaling | 8 | 4 | 0.92 | 0.00846 |

**Table S5.** OrthoFinder summary results.

| | *C. bombycina* | *C. holgerseni* | *Mel. bagoti* | *O. robustior* | *F. fusca* | *Myr. sabuleti* |
|---|---|---|---|---|---|---|
| **Number of genes** | 46,104 | 46,417 | 37,949 | 44,826 | 58,025 | 58,911 |
| **Number of genes in orthogroups** | 43,429 | 43,52 | 34,985 | 41,217 | 53,023 | 50,192 |
| **Number of unassigned genes** | 2,675 | 2,897 | 2,964 | 3,609 | 5,002 | 8,719 |
| **Percentage of genes in orthogroups** | 94.2 | 93.8 | 92.2 | 91.9 | 91.4 | 85.2 |
| **Percentage of unassigned genes** | 5.8 | 6.2 | 7.8 | 8.1 | 8.6 | 14.8 |
| **Number of orthogroups containing species** | 17,645 | 17,924 | 16,689 | 17,837 | 19,473 | 20,662 |
| **Percentage of orthogroups containing species** | 51.7 | 52.5 | 48.9 | 52.3 | 57.0 | 60.5 |
| **Number of species-specific orthogroups** | 712 | 859 | 960 | 1,637 | 1,748 | 3,053 |
| **Number of genes in species-specific orthogroups** | 2,315 | 2,594 | 2,904 | 5,099 | 6,400 | 10,333 |
| **Percentage of genes in species-specific orthogroups** | 5.0 | 5.6 | 7.7 | 11.4 | 11.0 | 17.5 |

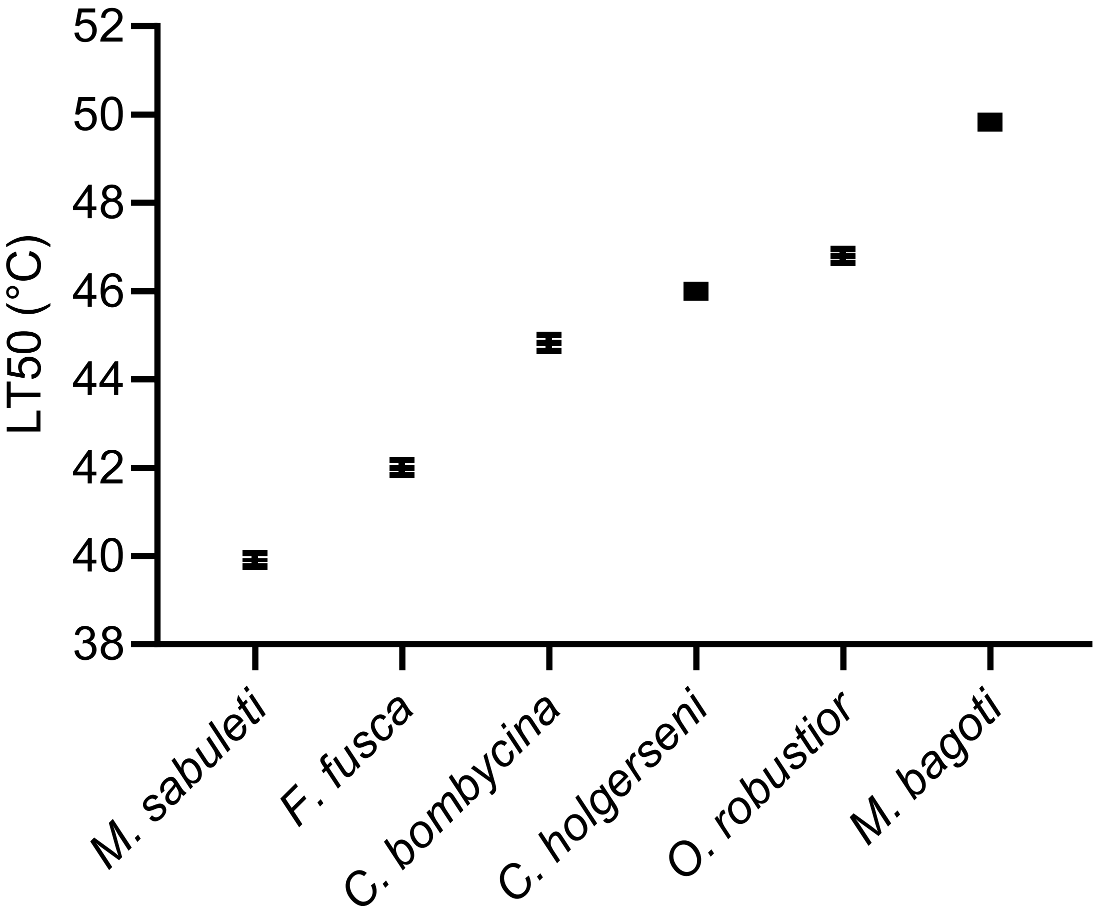


**Figure S1.** Median lethal temperature (LT50) after 3 hours exposure of studied species during the heat-stress treatment. LT50_3h and 95% confidence intervals were calculated using a simple logistic regression of death probability. A ratio test was used for pairwise comparisons between species. All species display significantly different values (*p* < 0.05).

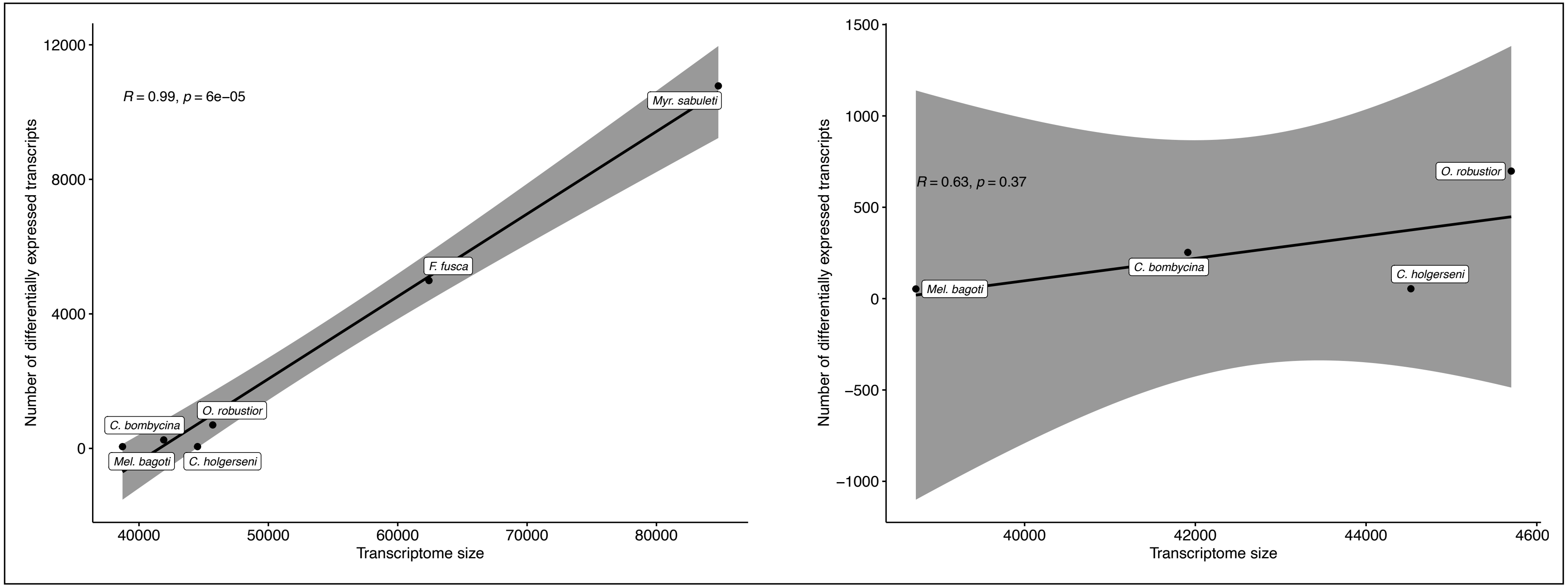


**Figure S2.** Correlation between total transcriptome size and the number of differentially expressed transcripts under heat stress. Left: all species included. Right: only desert ant species included. Correlation method Pearson, significance at $p < 0.05$.

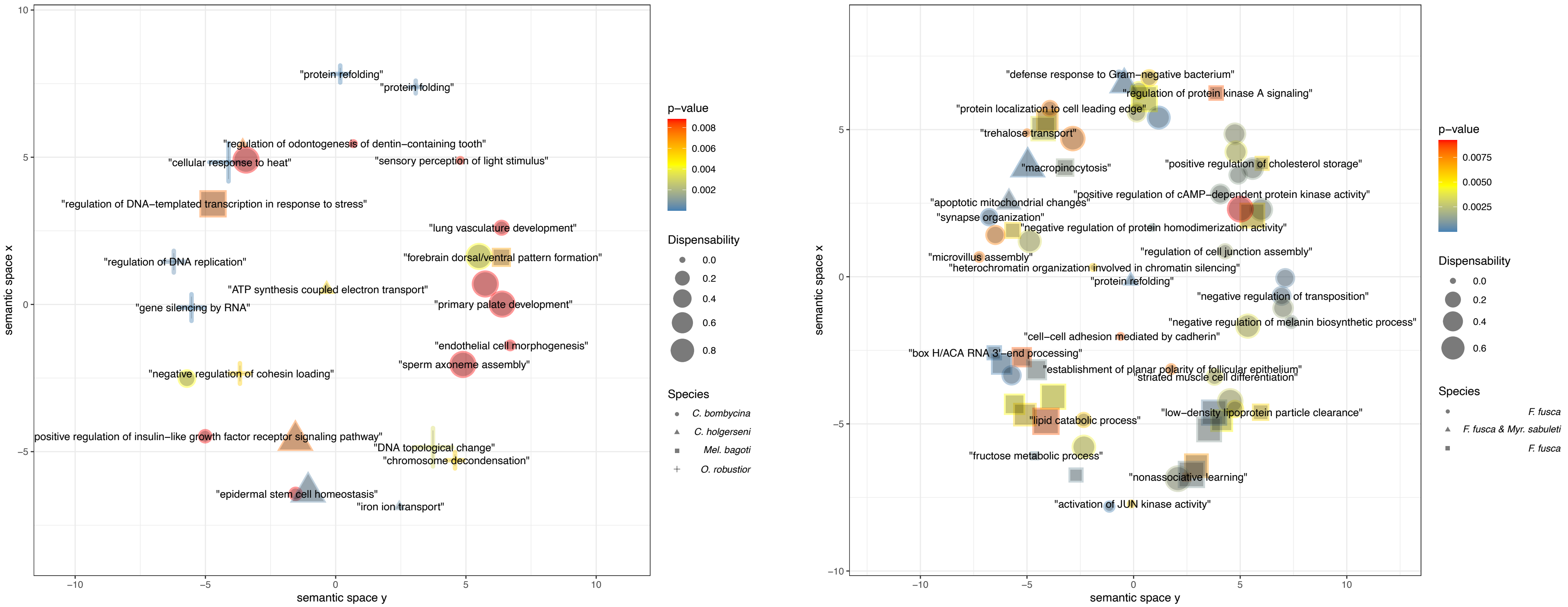


**Figure S3.** Summary of the enriched GO terms based on terms similarity as indicated by REVIGO. On the left is represented all GO terms enriched in desert ants, on the right enriched terms in temperate species. Colours indicate enrichment p-values, form size the dispensability of the term within the considered semantic space, and form shape shows in which species the term was significantly enriched.

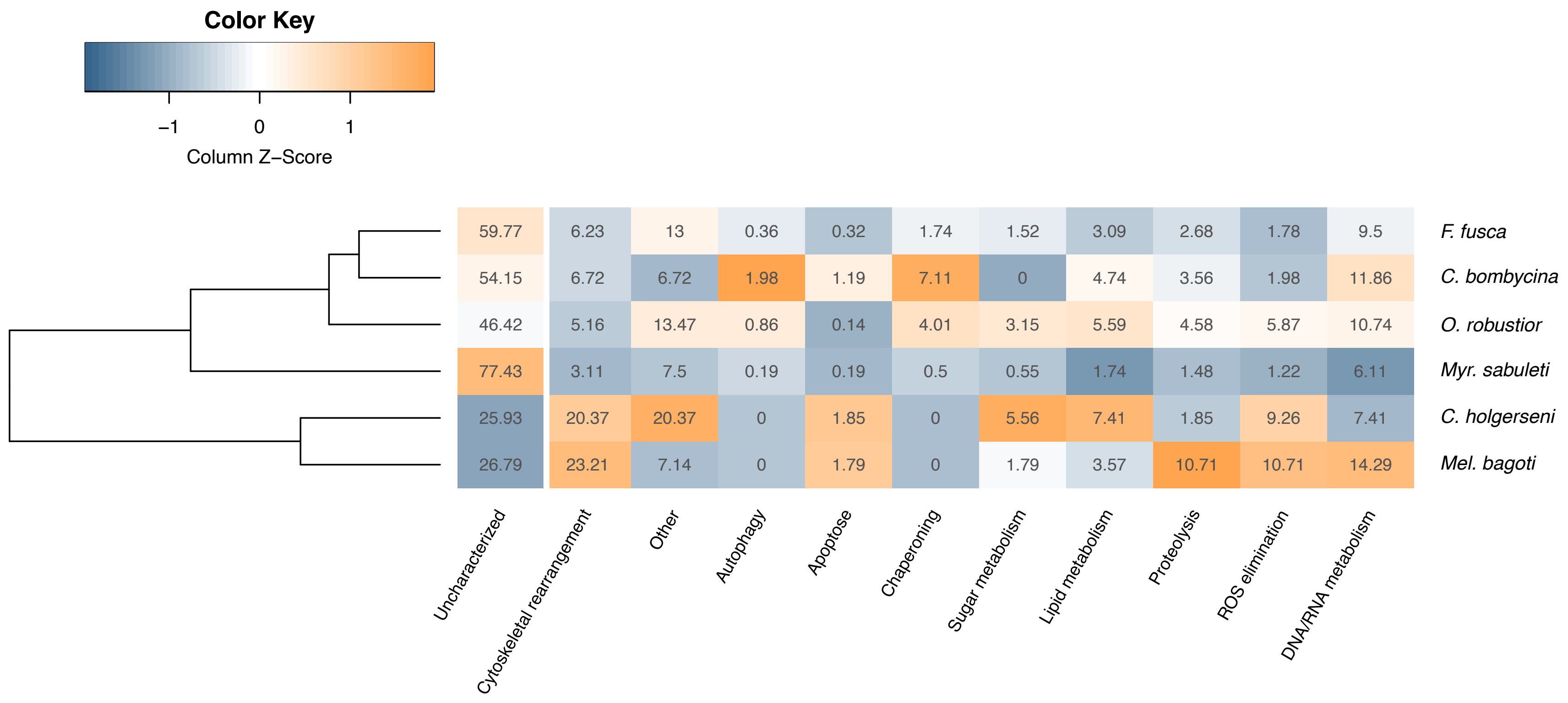


**Figure S4.** Relative proportion of differentially expressed transcripts involved in heat stress related functional categories. “Other” represent genes involved in any other biological function different from the ones detailed in the figure. “Uncharacterized” genes are shown for reference in the first column. Data were clustered according to their Euclidean distances and not according to species phylogenetic relationship. Colour is scaled by columns, i.e., for each column largest values are indicated in dark orange and smallest values in dark blue.

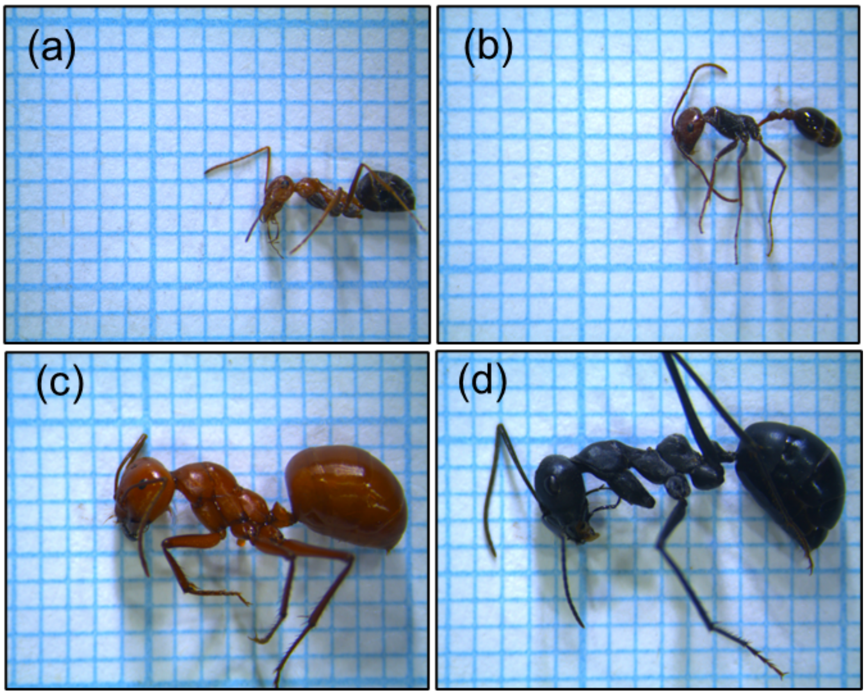


**Figure S5.** Workers of the desert ant species studied (a) *Cataglyphis bombycina*, (b) *Ocymyrmex robustior* (c) *Melophorus bagoti*, and (d) *Cataglyphis holgerseni*. The body mass of pictured workers was respectively: (a) 5.2 mg, (b) 4.6 mg (c) 59.8 mg and (d) 52.8 mg. Workers shown are representative of most foragers we observed.
